# The potential SARS-CoV-2 entry inhibitor

**DOI:** 10.1101/2020.03.26.009803

**Authors:** Jrhau Lung, Yu-Shih Lin, Yao-Hsu Yang, Yu-Lun Chou, Geng-He Chang, Ming-Shao Tsai, Cheng-Ming Hsu, Reming-Albert Yeh, Li-Hsin Shu, Yu-Ching Cheng, Hung Te Liu, Ching-Yuan Wu

## Abstract

Outbreak of coronavirus disease 2019 (COVID-19) occurred in Wuhan and has rapidly spread to almost all parts of world. In coronaviruses, the receptor binding domain (RBD) in the distal part of S1 subunit of SARS-CoV-2 spike protein can directly bind to angiotensin converting enzyme 2 (ACE2). RBD promote viral entry into the host cells and is an important therapeutic target. In this study, we discovered that theaflavin showed the lower idock score (idock score: −7.95 kcal/mol). To confirm the result, we discovered that theaflavin showed FullFitness score of −991.21 kcal/mol and estimated ΔG of −8.53 kcal/mol for the most favorable interaction with contact area of SARS-CoV-2 RBD by SwissDock service. Regarding contact modes, hydrophobic interactions contribute significantly in binding and additional hydrogen bonds were formed between theaflavin and Arg454, Phe456, Asn460, Cys480, Gln493, Asn501 and Val503 of SARS-CoV-2 RBD, near the direct contact area with ACE2. Our results suggest that theaflavin could be the candidate of SARS-CoV-2 entry inhibitor for further study.

## Introduction

In December 2019, an outbreak of coronavirus disease 2019 (COVID-19) occurred in Wuhan, China, induced by SARS-CoV-2. Human-to-human spread rapidly to the whole world triggering a global public health emergency (1–3). In coronaviruses, spike protein on the surface envelop of SARS-CoV-2 is responsible for promoting viral entry into the host cells (4). The spike protein is composed of two functional subunits: the S1 subunit that binds to the host cell receptors and the S2 subunit that mediates the fusion of the viral and host cellular membranes. The distal part of the S1 subunit contains the receptor binding domain (RBD) that directly binds to the peptidase domain of angiotensin converting enzyme 2 (ACE2)(5, 6). The directly contact area between RBD and ACE2 included several regions. Gln498, Thr500, Asn501 residues of the amino terminus (N) of RBD form several hydrogen bonds with the Tyr41, Gln42, Lys353, and Arg357 residues of ACE2. Lys417 and Tyr453 residues of RBD can interact with the Asp30 and His34 residues of ACE2, respectively. Gln474 in the carboxyl terminus (C) of RBD forms hydrogen bond with Gln24 of ACE2. In addition, Phe486 of RBD forms van der Waals forces with Met82 of ACE24 (Fig 1)(5). However, the sequence identity of the spike protein between SARS-CoV-2 and SARS-CoV is 76%, and major variation exists at the N-terminus encoding the RBD(7). Since RBD of spike protein is surface-exposed and promotes entry into host cells, it could be a potential target for therapy and vaccination using small molecules and neutralizing antibodies to treat COVID-19 (4, 8).

**Fig 1.**
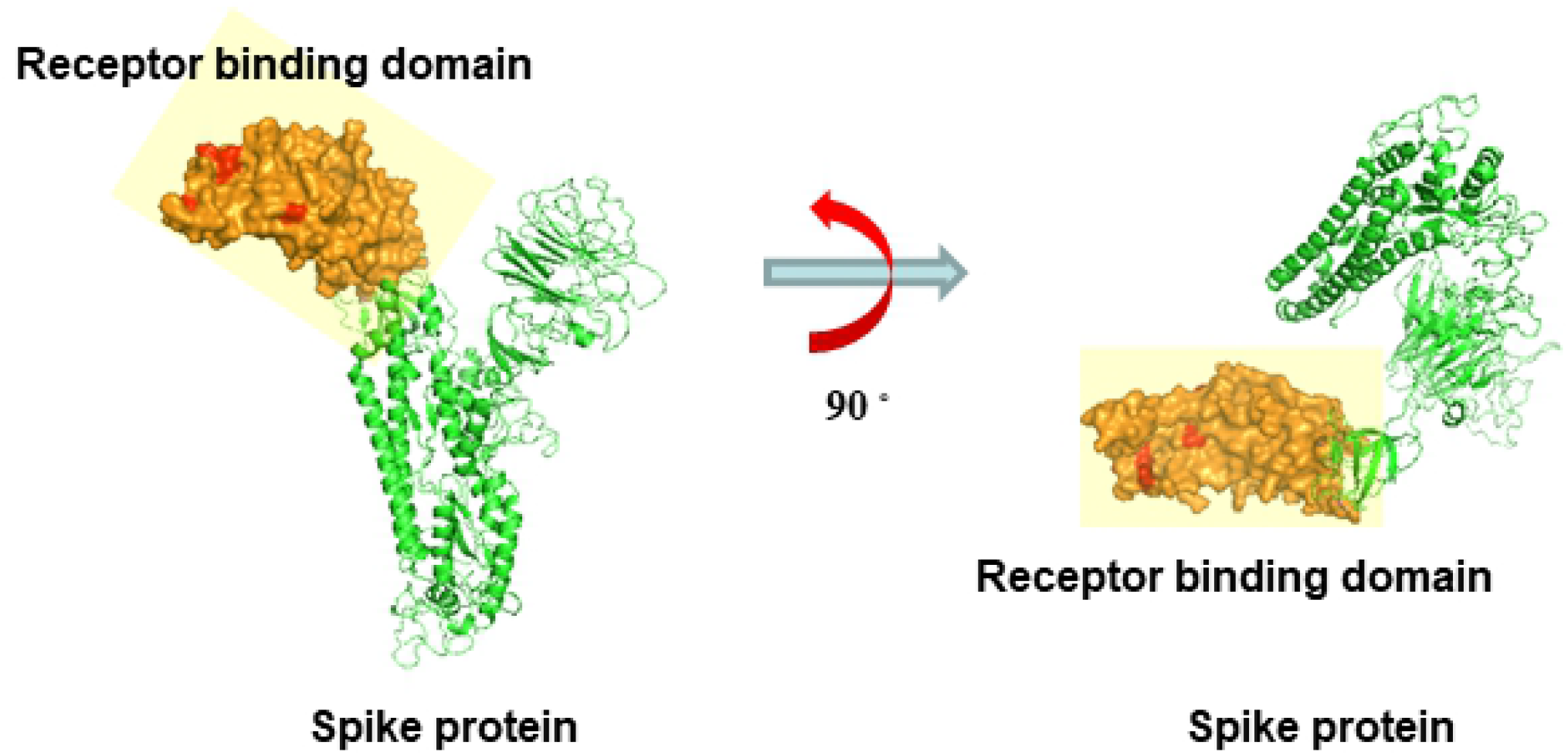
(**A**) A modeled structure of SARS-CoV-2 spike protein. Orange molecule represent crystallographic binding contact surface of RBD in SARS-CoV-2. Read molecule represent crystallographic directly binding amino acid residues of RBD in SARS-CoV-2. Grid box size for binding site is used by Modeller. The contact area and grid box size (light yellow) is showed for binding site.

In a recent study, we discovered that theaflavin possesses a potential chemical structure of anti-SARS-CoV-2 RNA-dependent RNA polymerase(9). Some Chinese medicinal compounds are also used for prophylaxis of SARS-CoV-2 infection. However, the actual mechanisms and efficiency of theaflavin for inhibiting SARS-CoV-2 entry into host cells are still unclear. Therefore, we used molecular docking to target the RBD of SARS-CoV-2, with the aim of screening these chemical structures of traditional Chinese medicinal compounds demonstrating commonly used against SARS-CoV and identifying alternative potential chemical structures for antiviral therapy.

## Methods and Materials

### Structure preparation

The 3D structure of spike protein of SARS-CoV-2 (NCBI Reference Sequence: YP_009724390.1) was generated based on homologous modeling using Modeller(10) incorporated within the UCSF Chimera(11) and SWISS-MODEL(12).

### Compound dataset collection

Eighty-three chemical structures from traditional Chinese medicinal compounds and their similar structures were retrieved from ZINC15 database.

### Molecular docking and virtual screening

We used two molecular docking methods for analysis. First, molecular docking and virtual screening was performed using idock download from Github (https://github.com/HongjianLi/idock) in a local linux machine. For each structure, nine docking posed were generated and the scores for the best docking poses of each structure were used for ranking by idock. Second, SwissDock server (Swiss Institute of Bioinformatics, University of Lausanne, Switzerland) was used for in silico prediction of the lowest free binding energy. The calculation was running online (accessible from http://www.swissdock.ch/) in the Internet browser(13). The grid box encompassed by Lys417 and Tyr453 of the middle of the bridge, Gln498, Thr500, Asn501 of the N-terminus, Gln474 of C-terminus and surrounding amino acids around contact surface of SARS-CoV-2 RBD was using for ligand docking and virtual screening (Fig 1).

## Result

Screening of these chemical structures, we discovered that theaflavin (ZINC3978446, Fig 2A) also has a lower idock score in the contact area of RBD in SARS-CoV-2 (−7.95 kcal/mol). The contact modes between theaflavin and RBD of SARS-CoV-2 with the lowest idocking scores are illustrated in Figure 1C. Regarding contact modes by idock, hydrophobic interactions contribute significantly in binding and additional hydrogen bond was found between theaflavin and SARS-CoV-2 RBD (Fig 2B).

**Fig 2.**
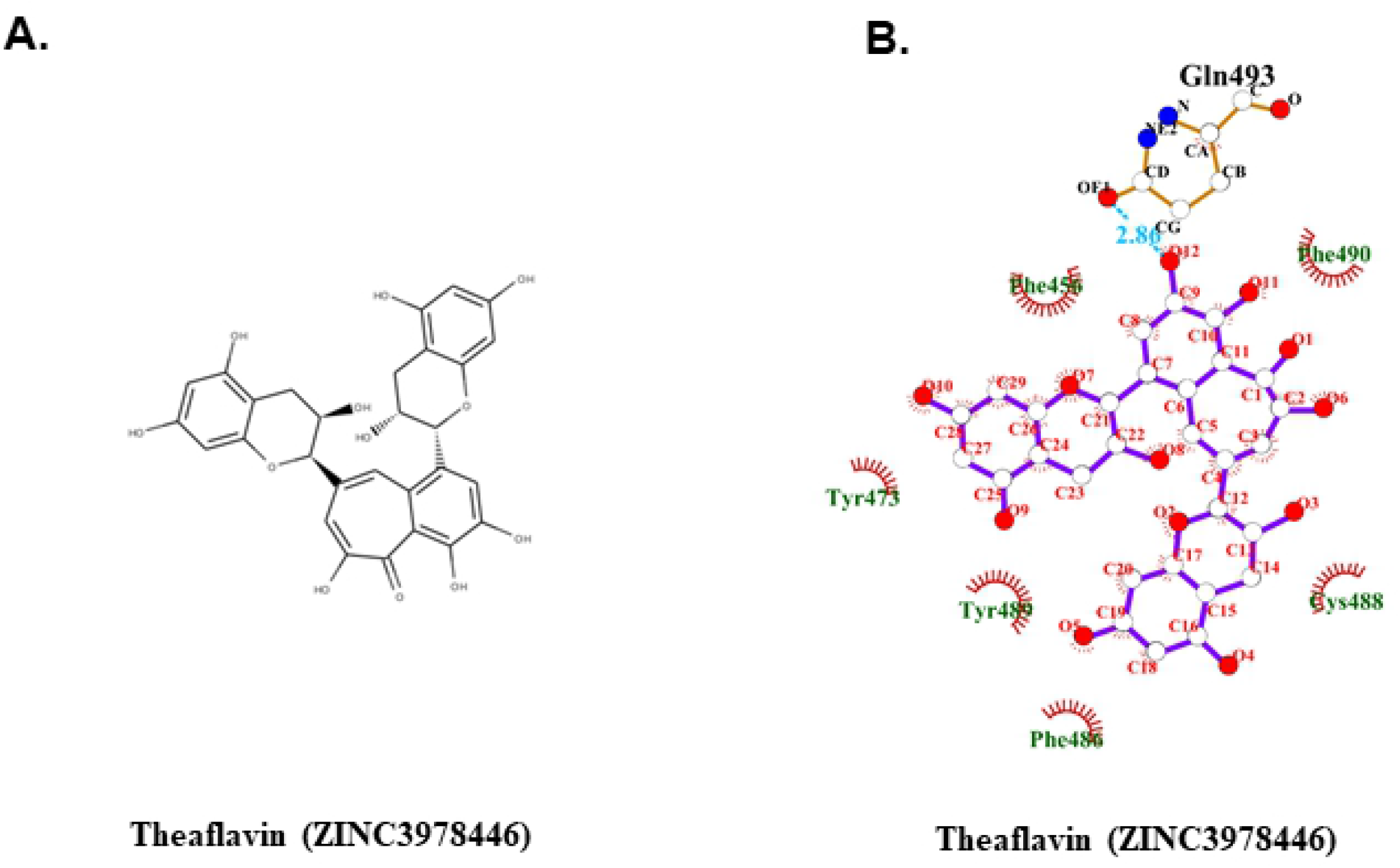
**(A)** The structure of theaflavin (ZINC3978446). (**B**) The contact model between theaflavin and SARS-CoV-2 RBD are showed in 2D interaction diagram by idock. Their relative distances between amino acid residues and catechin pentabenzoate or theaflavin digallte are analyzed and illustrated by LigPolt+. Carbon, oxygen, nitrogen, and fluoride molecules are marked as white, red, blue, and green circles, respectively. Covalent bonds in theaflavin and amino acid residues of RBD are labeled in purple and orange solid lines, respectively. The light blue dot lines label the distance (in Å) of hydrogen bonds formed between the functional moieties of theaflavin and amino acid residues. Hydrophobic interactions between theaflavin and RBD are depicted by the name of involving amino acid residues, which are labeled with dark green with dark red eyelashes pointing to the involved functional moiety of theaflavin.

Because theaflavin has the lower idock score in the contact area of SARS-CoV-2 RBD (−7.95 kcal/mol), we used the SwissDock server to confirm the result. Favorable binding modes were scored based on FullFitness scores and binding energy (Estimated ΔG (kcal/mol)) by the SwissDock server. The FullFitness scores and binding energy results and were obtained from the docking of theaflavin (ZINC3978446) into contact area of SARS-CoV-2 RBD. Theaflavin showed FullFitness scores of −991.21 kcal/mol and estimated ΔG of −8.53 kcal/mol for the most favorable interaction with contact area of SARS-CoV-2 RBD (Fig 3A). The 3D and 2D contact modes between contact area of SARS-CoV-2 RBD and theaflavin with the lowest binding energy are illustrated in Figure 3B, C and D. Regarding the contact modes by SwissDock server, hydrophobic interactions contribute significantly for binding (Fig 3D and E). The additional hydrogen bonds hydrogen bonds were formed between theaflavin and Arg454, Phe456, Asn460, Cys480, Gln493, Asn501 and Val503 of SARS-CoV-2 RBD, near the direct contact area with ACE2.

**Fig 3.**
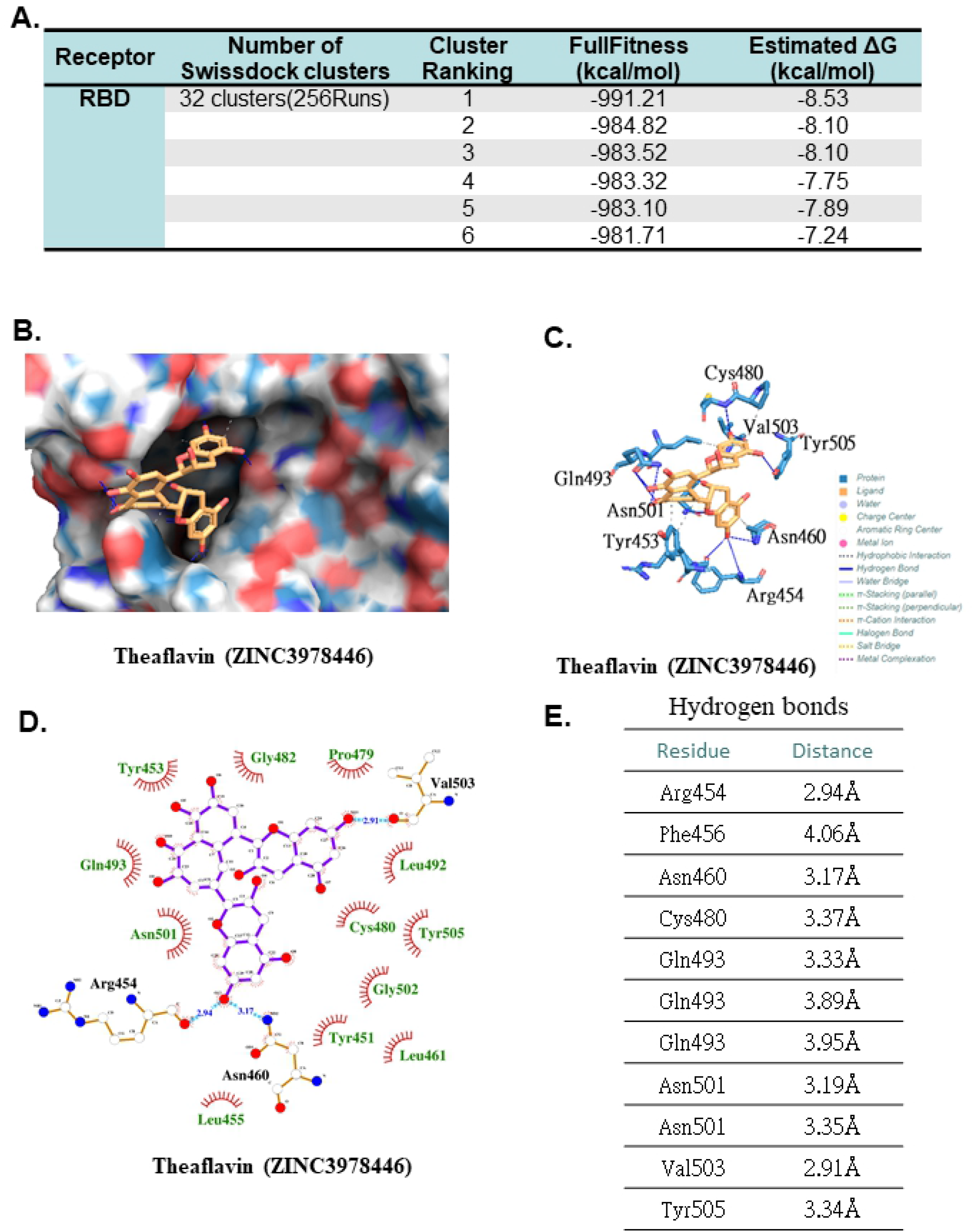
**(A)** Clustering results obtained from the docking of theaflavin into RBD by SwissDock service. **(B)** The molecules represent crystallographic and predicted pose for theaflavin in the pocket of RBD. **(C)** The hydrogen bonds interaction established by theaflavin with the closest residues of RBD are showed through Protein-Ligand Interaction Profiler (PLIP). **(D)** The contact model between theaflavin and SARS-CoV-2 RBD are showed in 2D interaction diagram by SwissDock. Their relative distances between amino acid residues and theaflavin are analyzed and illustrated by LigPolt+. Carbon, oxygen, nitrogen, and fluoride molecules are marked as white, red, blue, and green circles, respectively. Covalent bonds in theaflavin and amino acid residues of RBD are labeled in purple and orange solid lines, respectively. The light blue dot lines label the distance (in Å) of hydrogen bonds formed between the functional moieties of theaflavin and amino acid residues. Hydrophobic interactions between theaflavin and RBD are depicted by the name of involving amino acid residues, which are labeled with dark green with dark red eyelashes pointing to the involved functional moiety of theaflavin. **(E)** The table of hydrogen bonds between theaflavin and RBD by SwissDock.

## Discussion

The previous study reported that dihydrotanshinone I also had inhibitory effects against viral entry and replication in the MERS-CoV through the cell model (14). In addition, the recent study showed dihydrotanshinone I could inhibit viral entry through binding to the fusion cone of spike protein which is important for viral membrane fusion(15). The docking binding energy between dihydrotanshinone I and spike protein is −5.16 kcal/mol (15). In our previous study, we discovered that theaflavin (ZINC3978446) is a potential chemical structure of anti-SARS-CoV-2 RNA-dependent RNA polymerase(9). Moreover, we also found that theaflvin was able to dock in contact area in RBD of SARS-CoV-2. Theaflvin formed hydrophobic interactions and additional hydrogen bonds with the direct contact area of RBD. It is possible that theaflavin could ocuppy the contact area of RBD and block the interaction between ACE2 and RBD of SARS-CoV-2. Nevertheless, the mechanisms of viral entry blockage by dihydrotanshinone I and theaflavin could vary.

Our results suggest that theaflavin could be the candidate for prophylaxis or treatment of SARS-CoV-2 infection through RBD targeting. However, the exact in vivo effect remains unclear and further studies are necessary to confirm the effects of theaflavin against SARS-CoV-2 entry.

## Declarations

## Acknowledgement

This work was supported by grant CMRPG6H0162 from Chang Gung Memorial Hospital, and MOST 108-2320-B-182-021 from Ministry of Science and Technology to Dr. Ching Yuan Wu, and NMRPG6H0041 from Ministry of Science and Technology to Dr. Jrhau Lung. The authors thank the Health Information and Epidemiology Laboratory at the Chiayi Chang Gung Memorial Hospital for the comments and assistance in data analysis.

## Availability of data and materials

All data and materials are contained and described within the manuscript.

## Authors’ contributions

J. L. performed the experiments; C.Y.W. conceived the idea and designed experiments and wrote manuscript. Y.H.Y., L.H.S., G.H.C., M.S.T., Y.C.C., H.T.L., Y.S.L. and J.L. analyzed the data. Y.L.C., C.M.H. and R.A.Y. revised English writing of the manuscript. All authors reviewed and approved the final version.

## Competing interests

The authors declare that they have no competing interests.

## Consent for publication

Not applicable.

## Ethics approval and consent to participate

Not applicable.

